# Transfer of Xanthomonas campestris pv. arecae, and Xanthomonas campestris pv. musacearum to Xanthomonas vasicola (Vauterin) as Xanthomonas vasicola pv. arecae comb. nov., and Xanthomonas vasicola pv. musacearum comb. nov. and description of Xanthomonas vasicola pv. vasculorum pv. nov

**DOI:** 10.1101/571166

**Authors:** David J. Studholme, Emmanuel Wicker, Sadik Muzemil Abrare, Andrew Aspin, Adam Bogdanove, Kirk Broders, Zoe Dubrow, Murray Grant, Jeffrey B. Jones, Georgina Karamura, Jillian Lang, Jan Leach, George Mahuku, Gloria Valentine Nakato, Teresa Coutinho, Julian Smith, Carolee T. Bull

## LETTER TO THE EDITOR

Members of the genus *Xanthomonas*, within the gamma-Proteobacteria, collectively cause disease on more than 400 plant species (Hayward 1993), though some members are apparently non-pathogenic (Vauterin et al. 1996) and some have been isolated from clinical samples such as skin microbiota (Seité, Zelenkova, and Martin 2017). Historically, taxonomy of *Xanthomonas* was tied to the host of isolation (Starr 1981; Wernham 1948), with the genus being split into large numbers of species, each defined by this single phenotypic feature (Dye 1962). Subsequently, most of the species were transferred (lumped) into a single species, *X. campestris*, and designated as nomenspecies because the organisms could not be distinguished from one another by phenotypic and physiological tests (Lapage et al. 1992; Dye and Lelliott 1974). As a temporary solution, and to help to maintain a connection with the historical and plant pathological literature, these nomenspecies were designated as pathovars within *X. campestris*, each defined by host range or disease syndrome (Dye et al. 1980). More recently, DNA sequence comparisons and biochemical approaches revealed that some of the host ranges of pathovars of *X. campestris* were not correlated with inferred phylogenies (Parkinson et al. 2007, 2009; Rodriguez-R et al. 2012). There have been heroic advances to improve the taxonomy of the genus as a whole (Vauterin et al. 1990; Vauterin, Rademaker, and Swings 2000; Rademaker et al. 2005; Vauterin et al. 1995) and of individual taxa (da Gama et al. 2018; Constantin et al. 2016; Trébaol et al. 2000; Timilsina et al. 2019; Jones et al. 2004), based on phenotypic, chemotaxonomic and genotypic analyses. But in a number of taxa there remain unresolved issues.

The bacterial pathogen *X. campestris* pv. *musacearum* (Yirgou and Bradbury 1968) Dye 1978 presents a major threat to cultivation of banana and enset crops in central and eastern Africa, where it causes banana Xanthomonas wilt (BXW) and enset Xanthomonas wilt (EXW). Originally described as *X. musacearum* (Yirgou and Bradbury 1968), this pathogen was first isolated from enset and banana in the 1960s and early 1970s, respectively in Ethiopia (Yirgou and Bradbury 1968, 1974). Symptoms consistent with EXW were reported for Ethiopia as early as the 1930s (Castellani 1939). However, only in the 21^st^ century did the disease establish in the banana-growing areas of Burundi, Democratic Republic of Congo, Kenya, Rwanda, Tanzania and Uganda (Biruma et al. 2007; Tushemereirwe et al. 2004; Ndungo et al. 2006; Reeder et al. 2007; Carter et al. 2010). In this region around the Great Lakes of eastern and central Africa, BXW disease severely challenges the livelihoods and food security of millions (Blomme et al. 2017; Shimwela et al. 2016; Tinzaara et al. 2016; Blomme et al. 2013; Biruma et al. 2007; Nakato, Mahuku, and Coutinho 2018).

There is confusion in the literature about the taxonomy of this bacterium. Subsequent to its assignment to *X. campestris* (Young et al. 1978), molecular sequence and biochemical data indicated that this pathogen is more closely related to *X. vasicola* (Parkinson et al. 2007; Aritua et al. 2007) as detailed below. Thus, the first objective of this letter is to propose the transfer of *X. campestris* pv. *musacearum* (Yirgou and Bradbury 1968) Dye 1978 to *X. vasicola* Vauterin 1995. The second objective is to give a clear overview of the different evolutionary lineages that constitute the species *X. vasicola*, in the light of recent genomics analyses. Strains described as [*X. campestris* pv. zeae] (Qhobela, Claflin, and Nowell 1990; Coutinho and Wallis 1991) fall within a clade of *X. campestris* pv. *vasculorum* (Cobb 1894) Dye 1978 that belongs within the species *X. vasicola* Vauterin 1995. Furthermore, *X. campestris* pv. *arecae* (Rao and Mohan 1970) Dye 1978 is closely related to the type strain of *X. vasicola*, as judged by its *gyrB* sequence (Parkinson et al. 2009). In this manuscript, pathovar names that have no valid standing in nomenclature are presented with square brackets as is standard (Bull et al. 2012).

The species *X. vasicola* Vauterin 1995 was created to encompass *X. campestris* pv. *holcicola* (Elliott 1930) Dye 1978 and a subset of strains (not including the pathotype) of *X. campestris* pv. *vasculorum* (Cobb 1894) Dye 1978 (Young et al. 1978; Vauterin et al. 1995). Taxonomic studies revealed that *X. campestris* pv. *vasculorum* contained groups of strains that are clearly distinguishable from its pathotype strain by phenotypic and molecular traits, despite their shared host ranges (Vauterin et al. 1992; Péros et al. 1994; Dookun, Stead, and Autrey 2000; Stead 1989; Vauterin et al. 1995; Destéfano et al. 2003). Vauterin’s type-B strains are distinguished from type-A by SDS-PAGE of proteins, gas chromatography of fatty acid methyl esters and DNA-DNA hybridization (Yang et al. 1993). Type-A and type-B strains can also be distinguished by PCR-RFLP analysis (Destéfano et al. 2003). The pathotype strain of *X. campestris* pv. *vasculorum* belongs to type-A. Table 1 lists examples of *X. campestris* pv. *vasculorum* (Cobb 1894) Dye 1978 strains that were classified in one or more of those studies. Vauterin and colleagues assigned type-A strains to [*X. vasicola* pv. *vasculorum*], along with the pathotype, to *X. axonopodis* pv. *vasculorum* (Cobb) Vauterin, Hoste, Kersters & Swings and type-B (Vauterin et al. 1995). However, we note that this pathovar is invalid because of the lack of a formal proposal differentiating it from other pathovars (Young et al. 2004) and no designation of a pathotype strain.

**Table 1.**
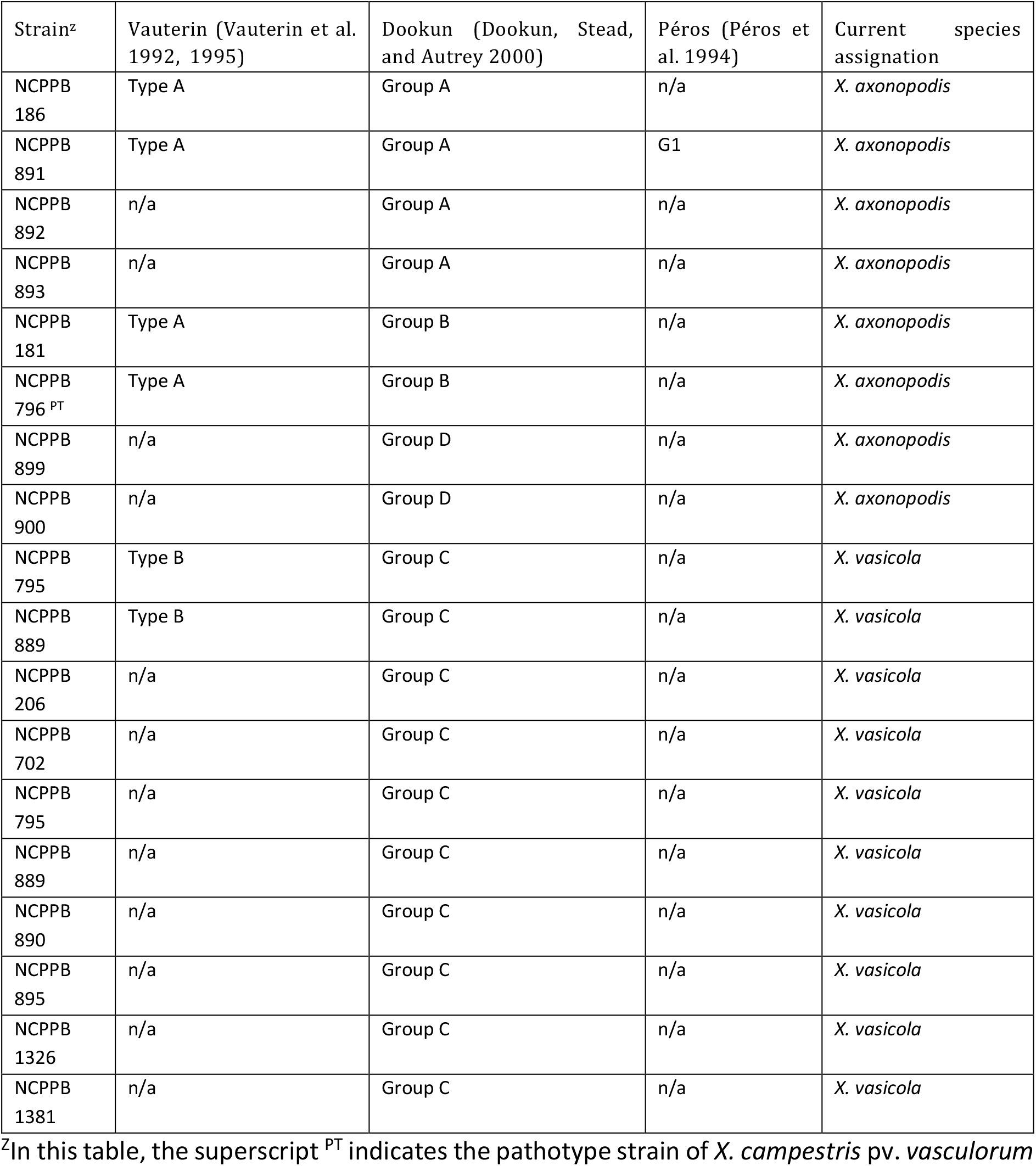
Classification of strains previously assigned to *X. campestris* pv. *vasculorum*.

Competing classifications and invalid names have led to the potentially confusing use of three different valid species names, *X. campestris, X. axonopodis* and *X. vasicola* to describe this group of bacteria in the literature. For example, various authors have referred to the single strain NCPPB 1326 as *X. campestris* pv. *vasculorum, X. axonopodis* pv. *vasculorum* (to which the strain clearly does not belong) or [*X. vasicola* pv. *vasculorum*] (Wasukira et al. 2014; Lewis Ivey, Tusiime, and Miller 2010; Qhobela, Claflin, and Nowell 1990; Qhobela and Claflin 1992). Type-B strains NCPPB 702, NCPPB 1326 and NCPPB 206 were erroneously described as *X. axonopodis* pv. *vasculorum* (Lewis Ivey, Tusiime, and Miller 2010) though they are clearly members of *X. vasicola*. However, we acknowledge that examples of mistakes such as these will not likely be resolved by transfer of the pathovars from *X. campestris* into *X. vasicola*.

A further source of confusion is the status of strains isolated from maize for which some authors use the invalid name [*X. campestris* pv. zeae] (Qhobela, Claflin, and Nowell 1990; Coutinho and Wallis 1991). Adding to the muddle, at least one strain of *X. campestris* pv. *vasculorum* (NCPPB 206) isolated from maize has the fatty-acid type characteristic of *X. vasicola* (Dookun, Stead, and Autrey 2000); consistent with this, on the basis of phylogenetic analysis of DNA sequence, this strain (NCPPB 206) clearly falls among strains assigned to Vauterin’s invalid [*X. vasicola* pv. *vasculorum*] (Wasukira et al. 2014). A useful nomenclature for this group has become more pressing since the recent outbreak of leaf streak on corn in the USA, caused by bacteria very closely related to strains previously described as [*X. campestris* pv. zeae]. One of these strains, NCPPB 4614 (=SAM119), has been suggested to be the eventual pathotype strain of *X. vasicola* pv. *vasculorum* though no valid proposal has been made (Lang et al. 2017; Korus et al. 2017). Although [*X. vasicola* pv. *vasculorum*] (Vauterin et al. 1995) is invalid, this name has come to be understood by the community to represent a meaningful biological reality; that is a set of *X. campestris* pv. *vasculorum* strains that are biochemically and phylogenetically similar to *X. vasicola*. Therefore, below we propose a formal description of *X. vasicola* pv. *vasculorum* pv. nov., which should be considered valid, to harmonize the formal nomenclature with that which is in use. Further, we therefore propose that [*X. vasicola* pv. *vasculorum*] group B, [*X. campestris* pv. zeae] and phylogenetically closely related strains isolated from sugarcane and maize be assigned into the newly described *X. vasicola* pv. *vasculorum* pv. nov.; this follows the previous suggestion (Lang et al. 2017) that strains classified to [*X. vasicola* pv. *vasculorum*] and [*X. campestris* pv. zeae] (Vauterin et al. 1995) are insufficiently distinct to warrant separate pathovars.

Vauterin et al. (1995) designated the pathotype strain of *X. vasicola* pv. *holcicola* (LMG 736, NCPPB 2417, ICMP 3103 and CFBP 2543) as the type strain of *X. vasicola*, although they did not use the pathovar epithet for the specific epithet of the species as is most appropriate to indicate this relationship. The natural host range of *X. vasicola* pv. *holcicola* includes the cereal crops millet and sorghum on which it causes bacterial leaf streak (Table 2). The host range of the strains that Vauterin et al. (1995) called [*X. vasicola* pv. *vasculorum*] is less well defined because in most of the relevant pre-1995 literature it is impossible to distinguish between type-A and type-B of *X. campestris* pv. *vasculorum* and therefore between *X. axonopodis* pv. *vasculorum* and strains belonging to *X. vasicola*. However, *X. campestris* pv. *vasculorum* type-B strains (that is, members of *X. vasicola*) have been isolated from sugarcane and maize and shown to infect these hosts on artificial inoculation (Vauterin et al. 1995; Karamura et al. 2015).

**Table 2.** Host ranges of the taxa discussed in this letter.

| Current taxon | Proposed taxon | Pathotype or Type strains | Additional strains in NCPPB known to be part of the newly proposed taxon | Natural hosts | Hosts by inoculation |
| --- | --- | --- | --- | --- | --- |
| *X. campestris* pv. *arecae* (Rao and Mohan 1970) Dye 1978 | *X. vasicola* pv. *arecae* pv. nov. | NCPPB 2649 = ICMP 5719 = LMG 533 | None | *Areca catechu* (Bradbury 1986; Kumar 1993, 1983) | *Cocos nucifera*, *Saccharum* sp. (Bradbury 1986) |
| *X. campestris* pv. *musacearum* (Yirgou and Bradbury 1968) Dye 1978 | *X. vasicola* pv. *musacearum* pv. nov | NCPPB 2005 = ATCC 49084 = CFBP 7123 = ICMP 2870 = LMG 785 | NCPPB 2251; NCPPB 4378; NCPPB 4379; NCPPB 4380; NCPPB 4381; NCPPB 4383; NCPPB 4384; NCPPB 4386; NCPPB 4387; NCPPB 4388; NCPPB 4389; NCPPB 4390; NCPPB 4391; NCPPB 4392; NCPPB 4393; NCPPB 4394; NCPPB 4395; NCPPB 4433; NCPPB 4434 | *Ensete ventricosum*, *Musa* sp. (Bradbury 1986), *Tripsacum* sp. (Unpublished observation, E. Wicker), | *Saccharum* sp., (Karamura et al. 2015), *Zea mays* (Karamura et al. 2015; Aritua et al. 2007) |
| [*Xanthomonas vasicola* pv. *zeae* Coutinho and Wallis 1990]<br>[*Xanthomonas vasicola* pv. *zeae* Qhobela et al 1990] | *X. vasicola* pv. *vasculorum* pv. nov. | NCPPB 4614 = SAM119 | None | *Zea mays* (Coutinho and Wallis 1991) | *Sorghum* sp. (Lang et al. 2017) |
| *X. vasicola* pv. *holcicola* (Elliott 1930) (Elliott 1930) Vauterin et al. 1995<br>(synonym of *X. campestris* pv. *holcicola*) | *X. vasicola* pv. *holcicola* (Elliott 1930) Vauterin et al. 1995 | NCPPB 2417 = CFBP 2543 = ICMP 3103 = LMG 736 | NCPPB 989; NCPPB 1060; NCPPB 1241; NCPPB 2417; NCPPB 2930; NCPPB 3162 | *Panicum miliaceum*, *Sorghum* spp., *Zea mays* (Bradbury 1986) | *Echinochloa frumentacea*, *Pennisetum typhoides*, *Setaria italica* (Bradbury 1986) |
| *X. campestris* pv. *vasculorum* type B = [*X. vasicola* pv. *vasculorum* (Vauterin et al., 1995)] | *X. vasicola* pv. *vasculorum* pv. nov. | NCPPB 4614 = SAM119 | NCPPB 206; NCPPB 702; NCPPB 795; NCPPB 889; NCPPB 890; NCPPB 895; NCPPB 1326; NCPPB 1381; NCPPB 4614 | *Saccharum* spp., *Zea mays*, *Eucalyptus grandis* (Coutinho et al. 2015; Bradbury 1986; Vauterin et al. 1995) | *Saccharum* spp., *Zea mays* (Karamura et al. 2015) |
| *Xanthomonas* sp. | *X. vasicola* Vauterin et al. 1995 | Not applicable | NCPPB 1394; NCPPB 1395; NCPPB 1396; NCPPB 902 | *Tripsacum laxum* (Mulder 1961), *Vetiveria zizanioides* (Kumar 1993, 1983) | Not known |

Previous studies suggested a close relationship between *X. campestris* pv. *musacearum* (Yirgou and Bradbury 1968) Dye 1978b and *X. vasicola* pv. *holcicola* (Elliott 1930) Vauterin et al. 1995 based on fatty acid methyl ester analysis, genomic fingerprinting using rep-PCR and partial nucleotide sequencing of the *gyrB* gene (Aritua et al. 2007; Parkinson et al. 2009). Draft or complete sequence assemblies are now available for more than a thousand *Xanthomonas* genomes, including those of type strains for most species and pathotypes for most pathovars. Genome-wide sequence data can offer some advantages, such as generally applicable threshold values for species delineation (Glaeser and Kämpfer 2015; Meier-Kolthoff et al. 2013; Meier-Kolthoff, Klenk, and Göker 2014; Richter and Rosselló-Móra 2009). Therefore, we further explored relationships among these organisms using whole genome sequences. We calculated pairwise average nucleotide identity (ANI) between *X. campestris* pv. *musacearum* and representative *Xanthomonas* strains, including all available species type strains and relevant pathotype strains. A representative subset of these pairwise ANI percentages is tabulated in Figure 1. This revealed that the pathotype strain (NCPPB 2005), of *X. campestris* pv. *musacearum* (Yirgou and Bradbury 1968) Dye 1978b shares 98.43 % ANI with the type strain of *X. vasicola* (NCPPB 2417) but only 87.27 % with the type strain of *X. campestris* (ATCC 33913). As expected, strains of *X. vasicola* pv. *holcicola* share high ANI (> 99.6 %) with the *X. vasicola* type strain, which is also the pathotype strain of *X. vasicola* pv. *holcicola* (Elliott 1930) Vauterin et al. 1995. Also as expected, strains of *X. campestris* pv. *vasculorum* previously called [*X. vasicola* pv. *vasculorum*] or [*X. campestris* pv. zeae], including the sequenced strain SAM119 (=NCPPB 4614) isolated from corn by T. Coutinho (Qhobela, Claflin, and Nowell 1990), share > 98.5 % ANI with the type strain of *X. vasicola*, supporting the need to transfer these strains to this species. Furthermore, unclassified strains NCPPB 902, NCPPB 1394, NCPPB 1395 and NCPPB 1396, from *Tripsacum laxum* (Mulder 1961) and the pathotype strain of *X. campestris* pv. *arecae* (Rao and Mohan 1970) Dye 1978 (NCPPB 2649) all share more than 98 % ANI with the type strain of *X. vasicola*, which places them unambiguously within *X. vasicola*. The next-nearest species to *X. vasicola* is *X. oryzae;* ANI between the respective type strains of these two species is 91.7%. It has been proposed that the boundary of a prokaryotic species can be delimited by 95 to 96% (Richter and Rosselló-Móra 2009). By this criterion, *X. campestris* pv. *arecae, X. campestris* pv. *musacearum* and strains from corn that are referred to by the invalid name *[Xanthomonas vasicola* pv. zeae] clearly fall within *X. vasicola* and outside *X. campestris*.

**Figure 1.**
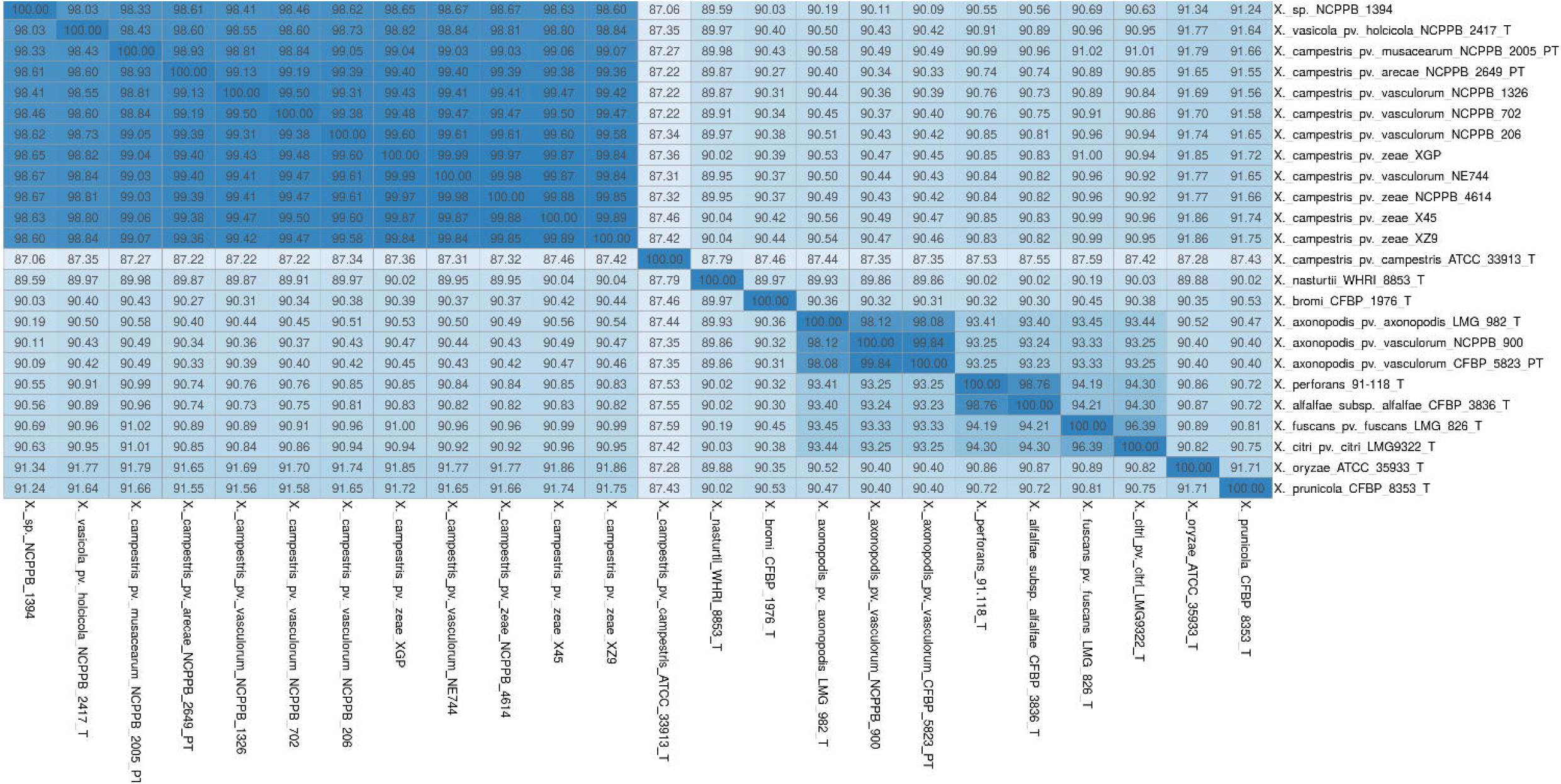
Average nucleotide identity (ANI) with type strains of *Xanthomonas* species. Genome sequence assemblies were obtained from GenBank and aligned against each other and ANI was calculated using the *dnadiff* function in MUMmer version 4 (Marçais et al. 2018). Accession numbers of the genome assemblies: GCA_000774005.1, GCA_000772705.1, GCA_000277875.1, GCA_000770355.1, GCA_000277995.1, GCA_000159795.2, GCA_000278035.1, GCA_003111865.1, GCA_002191965.1, GCA_002191955.1, GCA_003111905.1, GCA_003111825.1, GCA_000007145.1, GCA_001660815.1, GCA_002939755.1, GCA_001401595.1, GCA_002939725.1, GCA_000724905.2, GCA_000192045.3, GCA_000488955.1, GCA_001401605.1, GCA_002018575.1, GCA_000482445.1 and GCA_002846205.1 (Studholme et al. 2010; Wasukira et al. 2014, 2012; Lang et al. 2017; Sanko et al. 2018; da Silva et al. 2002; Vicente et al. 2017; Harrison and Studholme 2014; Potnis et al. 2011; Jacques et al. 2013).

The high ANI levels clearly delineate a genomospecies that includes the type strain *X. vasicola* NCPPB 2417. Nevertheless, despite the usefulness of ANI for delimiting species boundaries, it does not include any model of molecular evolution and thus is unsuited for phylogenetic reconstruction. Therefore, we used RaxML via the RealPhy pipeline (Bertels et al. 2014; Stamatakis, Ludwig, and Meier 2005) to elucidate phylogenetic relationships, using a maximum-likelihood method based on genome-wide sequencing data. This approach has the additional advantage of being based on sequence reads rather than on genome assemblies, where the latter may be of variable quality and completeness (Bertels et al. 2014).

Figure 2 depicts the phylogeny of *X. vasicola* based on RealPhy analysis of genome-wide sequence data. Pathovars *X. vasicola* pv. *holcicola* and *X. campestris* pv. *musacearum* are monophyletic, comprising well supported clades within the *X. vasicola* genomospecies. A third well supported clade includes the four *Xanthomonas* strains originating from the grass *Tripsacum laxum*. A fourth clade consists of mostly *X. campestris* pv. *vasculorum* strains isolated from sugarcane but also includes *X. campestris* pv. *vasculorum* strain NCPPB 206 isolated from maize and several strains from maize attributed to the invalid name [*X. campestris* pv. zeae]. This indicates that sequenced strains of [*X. campestris* pv. zeae] from corn (Sanko et al. 2018; Lang et al. 2017; Qhobela, Claflin, and Nowell 1990; Coutinho and Wallis 1991) are monophyletic and fall within the clade containing type-B strains of *X. campestris* pv. *vasculorum* (Figure 2). The single sequenced pathotype strain of *X. campestris* pv. *arecae* falls immediately adjacent to the *X. vasicola* clade containing strains from corn and *X. campestris* pv. *vasculorum* type B strains (Figure 2).

**Figure 2.**
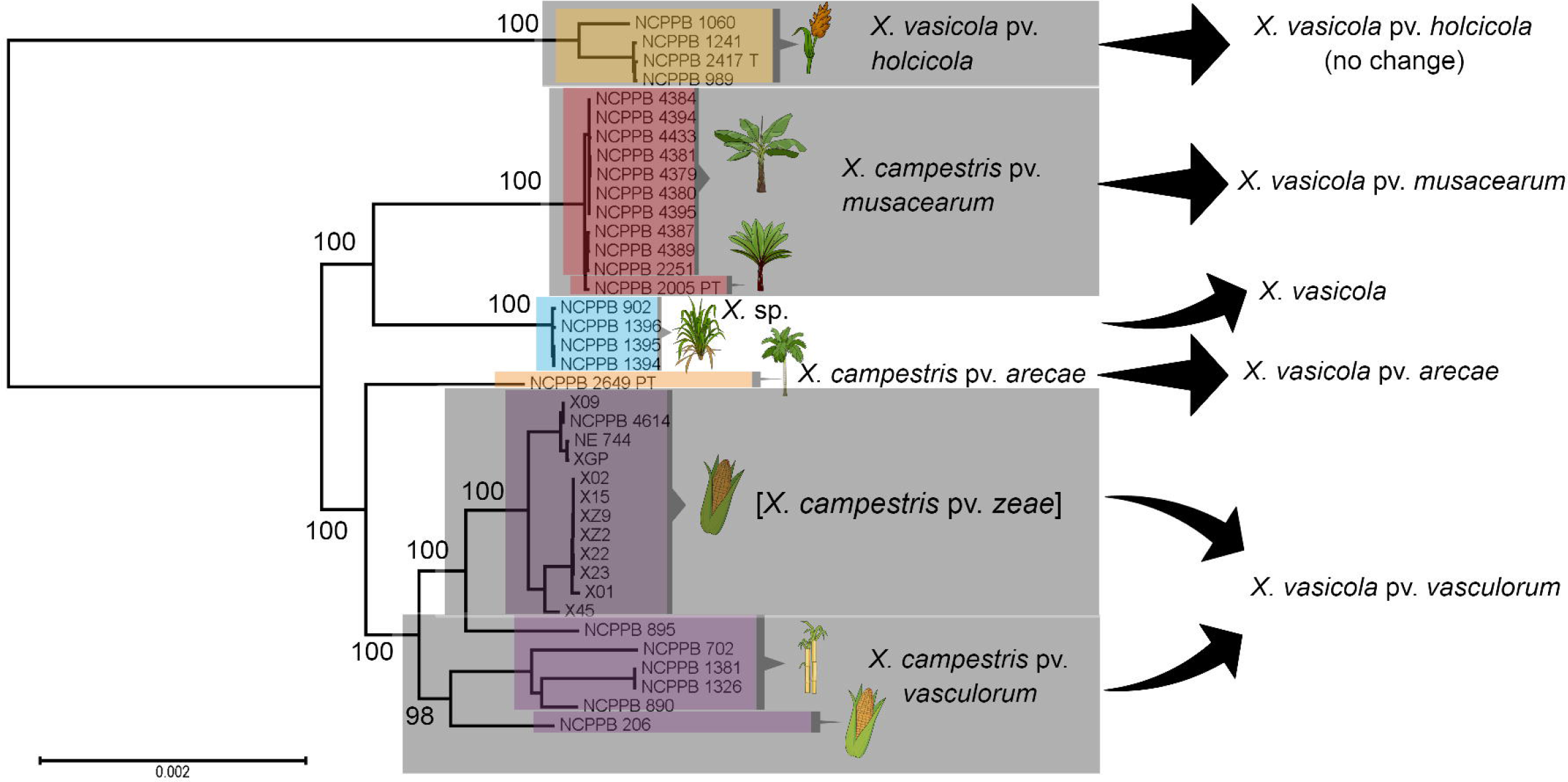
Maximum-likelihood phylogenetic tree based on genomic sequencing reads. The maximum likelihood tree was generated using RealPhy (Bertels et al. 2014) and RaxML (Stamatakis, Ludwig, and Meier 2005). Bootstrap values are expressed as percentages of 500 trials. Type and pathotype strains are indicated by ‘P’ and ‘PT’ respectively. Whole-genome shotgun sequence reads were obtained from the Sequence Read Archive (Leinonen, Sugawara, and Shumway 2011) via BioProjects PRJNA73853, PRJNA163305, PRJNA163307, PRJNA31213, PRJNA374510, PRJNA374557, PRJNA439013, PRJNA439327, PRJNA439328, PRJNA439329 and PRJNA449864 (Lang et al. 2017; Wasukira et al. 2014, 2012; Sanko et al. 2018).

Overall, our molecular sequence analyses strongly point to the existence of a phylogenetically coherent species, *X. vasicola* Vauterin 1995, that includes strains previously assigned to *X. campestris* pathovars *musacearum, arecae*, some strains of *X. campestris* pv. *vasculorum*, and strains collected from corn and *T. laxum* grass that have not been previously assigned to species nor pathovar. Here we propose that the pathovar *Xanthomonas vasicola* pv. *vasculorum* pv. nov. includes strains formerly classified as *X. campestris* pv. *vasculorum* but distinguishable from *X. axonopodis* pv. *vasculorum* (Cobb) Vauterin, Hoste, Kersters & Swings by protein SDS-PAGE, fatty acid methyl esterase (FAME) analysis and DNA hybridisation (Vauterin et al. 1992; Yang et al. 1993; Vauterin et al. 1995). Our analyses also support the transfer of *X. campestris* pv. arecae (Rao and Mohan 1970) Dye 1978 to *X. vasicola*. Although only a single genome of this pathovar has been sequenced, that genome belongs to the pathotype strain of the pathovar (Rao and Mohan 1970; Bull et al. 2010).

Our results are consistent with previous evidence for similarity between *X. campestris* pv. *musacearum* and strains of *X. vasicola*, based on FAME, genomic fingerprinting with rep-PCR and *gyrB* sequencing (Aritua et al. 2007; Parkinson et al. 2007). The formal species description for *X. vasicola* Vauterin 1995 states that this species can be clearly distinguished by its FAME profiles (Vauterin et al. 1995). Pathogenicity studies demonstrated phenotypic distinctiveness of *X. campestris* pv. *musacearum* (Yirgou and Bradbury 1968) Dye 1978 on banana; *X. campestris* pv. *musacearum* produces severe disease on this host whereas *X. vasicola* pv. *holcicola* NCPPB 2417 and *X. campestris* pv. *vasculorum* NCPPB 702 (which belongs to *X. vasicola*) induced no symptoms (Aritua et al. 2007). The species description (Vauterin et al. 1995) also states that *X. vasicola* is characterised by metabolic activity on the carbon substrates D-psicose and L-glutamic acid, and by a lack of metabolic activity on a range of carbon substrates (see below). We are not aware that these metabolic activities have been tested for *X. campestris* pv. *arecae, X. campestris* pv. *musacearum* and [*X. campestris* pv. zeae]; it is possible that the species description may need to be amended to accommodate any deviation from this definition among the repositioned pathovars.

Overall, it seems that the species *X. vasicola* (including *X. vasicola* pv. *holcicola, X. campestris* pv. *vasculorum* type-B strains, [*X. campestris* pv. zeae] strains, *X. campestris* pv. *arecae* and some strains isolated from *T. laxum*) is almost exclusively associated with monocot plants of the families Palmae and Gramineae. In this respect, it is similar to its closest sibling species *X. oryzae*, whose host range is limited to Gramineae (Bradbury 1986). The exception is a report of leaf blight and dieback in *Eucalyptus* caused by *X. vasicola* (Coutinho et al. 2015), remarkable given the phylogenetic distance between this dicot plant and the usual monocot hosts of *X. vasicola;* the infected South African plantation was in an area where sugarcane is grown.

In conclusion, analysis of available genome sequence data, combined with published pathogenicity and biochemical data, strongly support the transfer of the *X. campestris* pathovars *musacearum* and *arecae* to the species *X. vasicola* as, respectively, (i) *X. vasicola* pv. *musacearum* comb. nov. with NCPPB 2005 as the pathotype strain (being the type strain of *X. musacearum* and pathotype strain of *X. campestris* pv. *musacearum*) and (ii) *X. vasicola* pv. *arecae* comb. nov with NCPPB 2649 as the pathotype strain (being the type strain of *X. arecae* and pathotype strain of *X. campestris* pv. *arecae*). Strains NCPPB 206, NCPPB 702, NCPPB 795, NCPPB 890, NCPPB 895, NCPPB 1326, NCPPB 1381, and NCPPB 4614 form a phylogenetically and phenotypically coherent group with a distinctive host range causing symptoms on maize and sugarcane but not on banana (Aritua et al. 2007; Karamura et al. 2015) that falls within *X. vasicola* pv. *vasculorum* pv. nov. The strains isolated from *T. laxum* are also clearly within the phylogenetic bounds of *X. vasicola* but cannot be assigned to any pathovar and form a distinct clade. The previous proposal of [*X. vasicola* pv. *vasculorum*] was invalid due to the lack of a designated pathotype strain (Vauterin et al. 1995). We designate NCPPB 4614 as the pathotype strain for this pathovar, following the previous suggestion by Lang an colleagues (Lang et al. 2017). This strain was previously proposed as the pathotype of *X. vasicola* pv. *vasculorum* (Lang et al. 2017) and causes disease symptoms on maize and sugarcane (Lang et al. 2017) but not on banana (Supplementary Figure S1). Furthermore, given that strains from corn formerly described by the invalid name [*X. campestris* pv. zeae] are members of *X. vasicola* and have host ranges that cannot be distinguished from the pathotype strain of *X. vasicola* pv. *vasculorum*, we propose that these strains are members of this pathovar. Phylogenetic data support this as the corn strains represent a sub-clade within strains of *X. campestris* pv. *vasculorum* that fall within the emended *X. vasicola*.

## EMENDED DESCRIPTION OF *XANTHOMONAS VASICOLA* VAUTERIN ET. AL., 1995

The characteristics are as described for the genus and the species (Vauterin et al., 1995) extended with phylogenetic data from this study. The species can be clearly distinguished from other xanthomonads by MLSA and whole genome sequence analysis with members having more than 98 % ANI with the type strain. SDS-PAGE protein and FAME profiles have been shown to be distinguishing for some pathovars (Yang et al. 1993; Vauterin et al. 1992; Aritua et al. 2007), by the presence of metabolic activity on the carbon substrates D-psicose and L-glutamic acid, and by a lack of metabolic activity on the carbon substrates N-acetyl-D-galactosamine, L-arabinose, a-D-lactose, D-melibiose, P-methyl-D-glucoside, L-rhamnose, D-sorbitol, formic add, D-galactonic acid lactone, D-galacturonic acid, D-gluconic acid, D-glucuronic acid, p-hydroxyphenylacetic acid, a-ketovaleric acid, quinic acid, glucuronamide, L-asparagine, L-histidine, L-phenylalanine, urocanic acid, inosine, uridine, thymidine, DL-a-glycerol phosphate, glucose 1-phosphate, and glucose 6-phosphate. The G+C content is between 63.1 and 63.6 mol % as calculated from whole-genome sequence data. The type strain is *X. vasicola* pv. *holcicola* LMG 736 (= NCPPB 2417 = ICMP 3103 = CFBP 2543).

### *X. vasicola* pv. *holcicola* Vauterin et al., 1995

= *X. campestris* pv. *holcicola* (Elliott) Dye 1978.

Description is as presented by Vauterin et al. (1995). The pathovar is distinguished on the basis of phytopathogenic specialization. As shown here and elsewhere (Lang et al. 2017), the pathovar is distinct from other pathovars by MLSA and genome-wide sequence analysis. According to Bradbury (1986), gelatin and starch are hydrolysed by most isolates examined. The natural host range includes: *Panicum miliaceum, Sorghum* spp., *S. almum, S. bicolor (S. vulgare), S. caffrorum, S. durra, S. halepense, S. sudanense, S. technicum* (*S. bicolor* var. *technicus*), *Zea mays*. The artificial host range (by inoculation) includes *Echinochloa frumentacea, Pennisetum typhoides, Setaria italica*.

Pathotype strain: PDDCC 3103; NCPPB 2417.

### *X. vasicola* pv. *vasculorum* pv. nov

Description as for the species and this pathovar is distinguished on the basis of phytopathogenic specialization and includes the strains of the former taxon *X. campestris* pv. *vasculorum* type B and pathogens from corn. The pathovar is identified to species and distinguished from other pathovars by its *gyrB* gene sequence (Parkinson et al. 2009) and genome-wide sequence analysis. It is not known whether the strains being transferred to this taxon conform to the species description for metabolic activity. According to previously published work (Coutinho et al. 2015; Aritua et al. 2007; Karamura et al. 2015; Hayward 1962) the natural host range includes: *Saccharum* spp., *Zea mays, Eucalyptus grandis* and does not cause symptoms on banana (Supplementary Figure S1).

Pathotype strain: NCPPB 4614; SAM119.

### *X. vasicola* pv. *arecae* (Rao & Mohan) Dye 1978 comb. nov

= *X. campestris* pv. *arecae* (Rao & Mohan) Dye 1978.

Description as for the species and this pathovar is distinguished on the basis of phytopathogenic specialization. The pathovar is identified to species and distinguished from other pathovars by its *gyrB* gene sequence (Parkinson et al. 2009) and by genome-wide sequence analysis. According to Bradbury (1980) the natural host range includes: *Areca catechu* (areca nut). Bradbury (1986) reports the artificial host range to include: *Cocos nucifera* (coconut). Needle prick into sugar cane produced limited streaks, but the bacteria did multiply to some extent and could be re-isolated. Disease: leaf stripe. Long, narrow water-soaked lesions, becoming dark brown or black with age. It is not known if the strains being transferred to this taxon conform to the species description for metabolic activity.

Pathotype strain: NCPPB 2649; PDDCC 5791.

### *X. vasicola* pv. *musacearum* (Yirgou & Bradbury) Dye 1978 comb. nov

= *X. campestris* pv. *musacearum* (Yirgou & Bradbury) Dye 1978.

Description as for the species and this pathovar is identified to species and distinguished on the basis of phytopathogenic specialization and is distinct from other pathovars by its *gyrB* gene sequence (Parkinson et al. 2009) and genome-wide sequence analysis. Gelatin slowly liquefied, starch not hydrolysed. Growth quite rapid and very mucoid when cultured on Yeast-Peptone-Sucrose-agar based media for 48h at 28°C. According to Bradbury (1986), the natural hosts include: *Ensete ventricosum* (enset), *Musa* spp. (banana). Additional hosts by inoculation: *Saccharum* sp. (sugarcane), *Zea mays* (maize) and disease is exhibited as a bacterial wilt where leaves wilt and wither; yellowish bacterial masses are found in vascular tissue and parenchyma. It is not known if the strains being transferred to this taxon conform to the species description for metabolic activity.

Pathotype strain: NCPPB 2005; PDDCC 2870.

## Supporting information

Supplementary Figure S1. Pathogenicity on banana

---

**Supplementary Figure S1. Pathogenicity tests of *Xanthomonas vasicola* strains on *Musa acuminata* (AAA Group) ‘Grand Nain’.** Grand Nain banana plants in tissue culture 20 days post syringe inoculation at OD_600_ 0.2 with **A.** *X. campestris* pv. *musacearum* NCPPB 4433, **B.** 10 mM MgCl_2_, **C.** *X. campestris* pv. *vasculorum* SAM119 (=NCPPB 4614), **D.** *X. campestris* pv. *vasculorum* NCPPB 702.

## REFERENCES

Aritua, V., Parkinson, N., Thwaites, R., Heeney, J. V, Jones, D. R., Tushemereirwe, W., et al. 2007. Characterization of the *Xanthomonas* sp. causing wilt of enset and banana and its proposed reclassification as a strain of *X. vasicola*. Plant Pathol. 57:170–177.

Bertels, F., Silander, O. K., Pachkov, M., Rainey, P. B., and van Nimwegen, E. 2014. Automated reconstruction of whole-genome phylogenies from short-sequence reads. Mol. Biol. Evol. 31:1077–88.

Biruma, M., Pillay, M., Tripathi, L., Blomme, G., Abele, S., Mwangi, M., et al. 2007. Banana *Xanthomonas* wilt: A review of the disease, management strategies and future research directions. African J. Biotechnol. 6:953–962.

Blomme, G., Dita, M., Jacobsen, K., Pérez-Vicente, L., Molina, A., Ocimati, W., et al. 2017. Bacterial diseases of bananas and enset: current state of knowledge and integrated approaches towards sustainable management. Front. Plant Sci. 8:1290.

Blomme, G., Ploetz, R., Jones, D., De Langhe, E., Price, N., Gold, C., et al. 2013. A historical overview of the appearance and spread of *Musa* pests and pathogens on the African continent: highlighting the importance of clean *Musa* planting materials and quarantine measures. Ann. Appl. Biol. 162:4–26.

Bradbury, J. F. 1986. Guide to plant pathogenic bacteria. CAB International.

Bull, C. T., De Boer, S. H., Denny, T. P., Firrao, G., Fischer-Le Saux, M., Saddler, G. S., et al. 2012. List of new names of plant pathogenic bacteria (2008-2010). J. Plant Pathol. 94:21–27.

Bull, C. T., De Boer, S. H., Denny, T. P., Firrao, G., Saux, M. F.-L., Saddler, G. S., et al. 2010. Comprehensive List of Names of Plant Pathogenic Bacteria, 1980-2007. J. Plant Pathol. 92:551–592.

Carter, B. A., Reeder, R., Mgenzi, S. R., Kinyua, Z. M., Mbaka, J. N., Doyle, K., et al. 2010. Identification of *Xanthomonas vasicola* (formerly *X. campestris* pv. *musacearum*), causative organism of banana xanthomonas wilt, in Tanzania, Kenya and Burundi. Plant Pathol. 59:403.

Castellani, E. 1939. Su un marciume dell’Ensete. L’Agricoltura Colon. 33:297–300.

Cobb, N. A. 1894. Plant diseases and their remedies. Diseases of the sugarcane. Agric. Gaz. N. S. W. 4:808–833.

Constantin, E. C., Cleenwerck, I., Maes, M., Baeyen, S., Van Malderghem, C., De Vos, P., et al. 2016. Genetic characterization of strains named as *Xanthomonas axonopodis* pv. *dieffenbachiae* leads to a taxonomic revision of the *X. axonopodis* species complex. Plant Pathol. 65:792–806.

Coutinho, T. A., and Wallis, F. M. 1991. Bacterial Streak Disease of Maize (*Zea mays* L.) in South Africa. J. Phytopathol. 133:112–112.

Coutinho, T. A., van der Westhuizen, L., Roux, J., Mcfarlane, S. A., and Venter, S. N. 2015. Significant host jump of *Xanthomonas vasicola* from sugarcane to a *Eucalyptus grandis* clone in South Africa. Plant Pathol. 64:576–581.

Destéfano, S. A. L., Almeida, I. M. G., Rodrigues Neto, J., Ferreira, M., and Balani, D. M. 2003. Differentiation of *Xanthomonas* species pathogenic to sugarcane by PCR-RFLP analysis. Eur. J. Plant Pathol. 109:283–288.

Dookun, A., Stead, D. E., and Autrey, L. J. 2000. Variation among strains of Xanthomonas campestris pv. vasculorum from Mauritius and other countries based on fatty acid analysis. Syst. Appl. Microbiol. 23:148–55.

Dye, D., and Lelliott, R. 1974. Genus II. *Xanthomonas* Dowson 1939. In Bergey’s Manual of Determinative Bacteriology, eds. R. E. Buchanan and N. E. Gibbons. Williams and Wilkins Co., Baltimore, U.S.A., p. 243–249.

Dye, D. W. 1962. The inadequacy of the usual determinative tests for the identification of *Xanthomonas* spp. New Zeal. J. Sci. 5:393–416.

Dye, D. W., Bradbury, J. F., Goto, M., Hayward, A. C., Lelliott, R. A., and Schroth, M. N. 1980. International standards for naming pathovars of phytopathogenic bacteria and a list of pathovar names and pathotype strains. Rev. Plant Pathol. 59:153–168.

Elliott, C. 1930. Bacterial streak disease of Sorghums. J. Agric. Res. 40:963–976 pp.

da Gama, M. A. S., Mariano, R. de L. R., da Silva Júnior, W. J., de Farias, A. R. G., Barbosa, M. A. G., Ferreira, M. Á. da S. V., et al. 2018. Taxonomic Repositioning of *Xanthomonas campestris* pv. *viticola* (Nayudu 1972) Dye 1978 as *Xanthomonas citri* pv. *viticola* (Nayudu 1972) Dye 1978 comb. nov. and Emendation of the Description of *Xanthomonas citri* pv. <i>anac. Phytopathology. 108:1143–1153.

Glaeser, S. P., and Kämpfer, P. 2015. Multilocus sequence analysis (MLSA) in prokaryotic taxonomy. Syst. Appl. Microbiol. 38:237–45.

Harrison, J., and Studholme, D. J. 2014. Draft genome sequence of *Xanthomonas axonopodis* pathovar *vasculorum* NCPPB 900. FEMS Microbiol. Lett. 360:113–6.

Hayward, A. C. 1962. Studies on Bacterial Pathogens of Sugar Cane. Mauritius Sugar Industry Research Institute.

Hayward, A. C. 1993. The hosts of *Xanthomonas*. In Xanthomonas, eds. J G Swings and E L Civerolo. Dordrecht: Springer Netherlands, p. 1–119.

Jacques, M. A., Bolot, S., Charbit, E., Darrasse, A., Briand, M., Arlat, M., et al. 2013. High-quality draft genome sequence of *Xanthomonas alfalfae* subsp. *alfalfae* strain CFBP 3836. Genome Announc. 1:e01035–13–e01035–13.

Jones, J. B., Lacy, G. H., Bouzar, H., Stall, R. E., and Schaad, N. W. 2004. Reclassification of the Xanthomonads Associated with Bacterial Spot Disease of Tomato and Pepper. Syst. Appl. Microbiol. 27:755–762.

Karamura, G., Smith, J., Studholme, D., Kubiriba, J., and Karamura, E. 2015. Comparative pathogenicity studies of the *Xanthomonas vasicola* species on maize, sugarcane and banana. African J. Plant Sci. 9:385–400.

Korus, K., Lang, J. M., Adesemoye, A. O., Block, C. C., Pal, N., Leach, J. E., et al. 2017. First Report of *Xanthomonas vasicola* causing bacterial leaf streak on corn in the United States. Plant Dis. 101:1030.

Kumar, S. N. S. 1983. Epidemiology of bacterial leaf stripe disease of arecanut palm. Trop. Pest Manag. 29:249–252.

Kumar, S. N. S. 1993. Perpetuation and host range of *Xanthomonas campestris* pv. *arecae* incitant of bacterial leaf stripe disease of arecanut palm. Adv. Hortic. For. 3:99–103.

Lang, J. M., DuCharme, E., Ibarra Caballero, J., Luna, E., Hartman, T., Ortiz-Castro, M., et al. 2017. Detection and characterization of *Xanthomonas vasicola* pv. *vasculorum* (Cobb 1894) comb. nov. causing bacterial leaf streak of corn in the United States. Phytopathology. 107:1312–1321.

Lapage, S. P., Sneath, P. H. A., Lessel, E. F., Skerman, V. B. D., Seeliger, H. P. R., and Clark, W. A., eds. 1992. International Code of Nomenclature of Bacteria: Bacteriological Code, 1990 Revision. Washington (DC).

Leinonen, R., Sugawara, H., and Shumway, M. 2011. The Sequence Read Archive. Nucleic Acids Res. 39:D19–D21.

Lewis Ivey, M. L., Tusiime, G., and Miller, S. A. 2010. A polymerase chain reaction assay for the detection of *Xanthomonas campestris* pv. *musacearum* in banana. Plant Dis. 94:109–114.

Marçais, G., Delcher, A. L., Phillippy, A. M., Coston, R., Salzberg, S. L., and Zimin, A. 2018. MUMmer4: A fast and versatile genome alignment system ed. Aaron E. Darling. PLOS Comput. Biol. 14:e1005944.

Meier-Kolthoff, J. P., Auch, A. F., Klenk, H.-P., and Göker, M. 2013. Genome sequence-based species delimitation with confidence intervals and improved distance functions. BMC Bioinformatics. 14:60.

Meier-Kolthoff, J. P., Klenk, H.-P., and Göker, M. 2014. Taxonomic use of DNA G+C content and DNA-DNA hybridization in the genomic age. Int. J. Syst. Evol. Microbiol. 64:352–6.

Mulder, D. 1961. A Bacterial Disease of Guatemala Grass. Tea Q. 32:143–144.

Nakato, V., Mahuku, G., and Coutinho, T. 2018. *Xanthomonas campestris* pv. *musacearum:* a major constraint to banana, plantain and enset production in central and east Africa over the past decade. Mol. Plant Pathol. 19:525–536.

Ndungo, V., Eden-Green, S., Blomme, G., Crozier, J., and Smith, J. J. 2006. Presence of banana xanthomonas wilt (*Xanthomonas campestris* pv. *musacearum*) in the Democratic Republic of Congo (DRC). Plant Pathol. 55:294.

Parkinson, N., Aritua, V., Heeney, J., Cowie, C., Bew, J., and Stead, D. 2007. Phylogenetic analysis of *Xanthomonas* species by comparison of partial gyrase B gene sequences. Int. J. Syst. Evol. Microbiol. 57:2881–7.

Parkinson, N., Cowie, C., Heeney, J., and Stead, D. 2009. Phylogenetic structure of *Xanthomonas* determined by comparison of *gyrB* sequences. Int. J. Syst. Evol. Microbiol. 59:264–74.

Péros, J. P., Girard, J. C., Lombard, H., Janse, J. D., and Berthier, Y. 1994. Variability of *Xanthomonas campestris* pv. *vasculorum* from sugarcane and other Gramineae in Reunion Island. Characterization of a different xanthomonad. J. Phytopathol. 142:177–188.

Potnis, N., Krasileva, K., Chow, V., Almeida, N. F., Patil, P. B., Ryan, R. P., et al. 2011. Comparative genomics reveals diversity among xanthomonads infecting tomato and pepper. BMC Genomics. 12:146.

Qhobela, M., and Claflin, L. E. 1992. Eastern and southern African strains of *Xanthomonas campestris* pv. *vasculorum* are distinguishable by restriction fragment length polymorphism of DNA and polyacrylamide gel electrophoresis of membrane proteins. Plant Pathol. 41:113–121.

Qhobela, M., Claflin, L. E., and Nowell, D. C. 1990. Evidence that *Xanthomonas campestris* pv. *zeae* can be distinguished from other pathovars capable of infecting maize by restriction fragment length polymorphism of genomic DNA. Can. J. Plant Pathol. 12:183–186.

Rademaker, J. L. W., Louws, F. J., Schultz, M. H., Rossbach, U., Vauterin, L., Swings, J., et al. 2005. A comprehensive species to strain taxonomic framework for *Xanthomonas*. Phytopathology. 95:1098–111.

Rao, Y. P., and Mohan, S. K. 1970. A new bacterial leaf stripe disease of arecanut (*Areca catechu*) in Mysore State. Indian Phytopathol. 23:702–704.

Reeder, R. H., Muhinyuza, J. B., Opolot, O., Aritua, V., Crozier, J., and Smith, J. 2007. Presence of banana bacterial wilt (*Xanthomonas campestris* pv. *musacearum*) in Rwanda. Plant Pathol. 56:1038.

Richter, M., and Rosselló-Móra, R. 2009. Shifting the genomic gold standard for the prokaryotic species definition. Proc. Natl. Acad. Sci. U. S. A. 106:19126–31.

Rodriguez-R, L. M., Grajales, A., Arrieta-Ortiz, M., Salazar, C., Restrepo, S., and Bernal, A. 2012. Genomes-based phylogeny of the genus *Xanthomonas*. BMC Microbiol. 12:43.

Sanko, T. J., Kraemer, A. S., Niemann, N., Gupta, A. K., Flett, B. C., Mienie, C., et al. 2018. Draft genome assemblages of 10 *Xanthomonas vasicola* pv. *zeae* strains, pathogens causing leaf streak disease of maize in South Africa. Genome Announc. 6:1–2.

Seité, S., Zelenkova, H., and Martin, R. 2017. Clinical efficacy of emollients in atopic dermatitis patients - relationship with the skin microbiota modification. Clin. Cosmet. Investig. Dermatol. 10:25–33.

Shimwela, M. M., Ploetz, R. C., Beed, F. D., Jones, J. B., Blackburn, J. K., Mkulila, S. I., et al. 2016. Banana xanthomonas wilt continues to spread in Tanzania despite an intensive symptomatic plant removal campaign: an impending socio-economic and ecological disaster. Food Secur. 8:939–951.

da Silva, A. C. R., Ferro, J. A., Reinach, F. C., Farah, C. S., Furlan, L. R., Quaggio, R. B., et al. 2002. Comparison of the genomes of two *Xanthomonas* pathogens with differing host specificities. Nature. 417:459–63.

Stamatakis, A., Ludwig, T., and Meier, H. 2005. RAxML-III: a fast program for maximum likelihood-based inference of large phylogenetic trees. Bioinformatics. 21:456–63.

Starr, M. P. 1981. The genus *Xanthomonas*. In The Prokaryotes, eds. Mortimer P. Starr, Heinz Stolp, Hans G. Trüper, Albert Balows, and Hans G. Schlegel. Berlin, Heidelberg: Springer Berlin Heidelberg, p. 742–763.

Stead, D. E. 1989. Grouping of *Xanthomonas campestris* pathovars of cereals and grasses by fatty acid profiling. EPPO Bull. 19:57–68.

Studholme, D. J., Kemen, E., MacLean, D., Schornack, S., Aritua, V., Thwaites, R., et al. 2010. Genome-wide sequencing data reveals virulence factors implicated in banana *Xanthomonas* wilt. FEMS Microbiol. Lett. 310:182–192.

Timilsina, S., Kara, S., Jacques, M. A., Potnis, N., Minsavage, G. V., Vallad, G. E., et al. 2019. Reclassification of *Xanthomonas gardneri* (ex Šutič 1957) Jones et al. 2006 as a later heterotypic synonym of *Xanthomonas cynarae* Trébaol et al. 2000 and description of *X. cynarae* pv. *cynarae* and *X. cynarae* pv.<i> gardner. Int. J. Syst. Evol. Microbiol. 69:343–349.

Tinzaara, W., Karamura, E. B., Kubiriba, J., Ochola, D., Ocimati, W., Blomme, G., et al. 2016. The banana *Xanthomonas* wilt epidemic in east and central Africa: current research and development efforts. Acta Hortic.:267–274.

Trébaol, G., Gardan, L., Manceau, C., Tanguy, J. L., Tirilly, Y., and Boury, S. 2000. Genomic and phenotypic characterization of *Xanthomonas cynarae* sp. nov., a new species that causes bacterial bract spot of artichoke (*Cynara scolymus* L.). Int. J. Syst. Evol. Microbiol. 50 Pt 4:1471–8.

Tushemereirwe, W., Kangire, A., Ssekiwoko, F., Offord, L. C., Crozier, J., Boa, E., et al. 2004. First report of *Xanthomonas campestris* pv. *musacearum* on banana in Uganda. Plant Pathol. 53:802–802.

Vauterin, L., Hoste, B., Kersters, K., and Swings, J. 1995. Reclassification of *Xanthomonas*. Int. J. Syst. Bacteriol. 45:472–489.

Vauterin, L., Rademaker, J., and Swings, J. 2000. Synopsis on the taxonomy of the genus *Xanthomonas*. Phytopathology. 90:677–82.

Vauterin, L., Swings, J., Kersters, K., Gillis, M., Mew, T. W., Schroth, M. N., et al. 1990. Towards an Improved Taxonomy of *Xanthomonas*. Int. J. Syst. Bacteriol. 40:312–316.

Vauterin, L., Yang, P., Alvarez, A., Takikawa, Y., Roth, D. A., Vidaver, A. K., et al. 1996. Identification of non-pathogenic *Xanthomonas* strains associated with plants. Syst. Appl. Microbiol. 19:96–105.

Vauterin, L., Yang, P., Hoste, B., Pot, B., Swings, J., and Kersters, K. 1992. Taxonomy of xanthomonads from cereals and grasses based on SDS-PAGE of proteins, fatty acid analysis and DNA hybridization. J. Gen. Microbiol. 138:1467–1477.

Vicente, J. G., Rothwell, S., Holub, E. B., and Studholme, D. J. 2017. Pathogenic, phenotypic and molecular characterisation of *Xanthomonas nasturtii* sp. nov. and *Xanthomonas floridensis* sp. nov., new species of *Xanthomonas* associated with watercress production in Florida. Int. J. Syst. Evol. Microbiol. 67:3645–3654.

Wasukira, A., Coulter, M., Al-Sowayeh, N., Thwaites, R., Paszkiewicz, K., Kubiriba, J., et al. 2014. Genome sequencing of *Xanthomonas vasicola* pathovar *vasculorum* reveals variation in plasmids and genes encoding lipopolysaccharide synthesis, type-IV pilus and type-III secretion effectors. Pathog. (Basel, Switzerland). 3:211–37.

Wasukira, A., Tayebwa, J., Thwaites, R., Paszkiewicz, K., Aritua, V., Kubiriba, J., et al. 2012. Genome-wide sequencing reveals two major sub-lineages in the genetically monomorphic pathogen *Xanthomonas campestris* pathovar *musacearum*. Genes (Basel). 3:361–377.

Wernham, C. C. 1948. The species value of pathogenicity in the genus *Xanthomonas*. Am. Phytopathol. Soc. 38:283–291.

Yang, P., Vauterin, L., Vancanneyt, M., Swings, J., and Kersters, K. 1993. Application of Fatty Acid Methyl Esters for the Taxonomic Analysis of the Genus *Xanthomonas*. Syst. Appl. Microbiol. 16:47–71.

Yirgou, D., and Bradbury, J. F. 1974. A note on wilt of banana caused by the enset wilt organism *Xanthomonas musacearum*. East African Agric. For. J. 40:111–114.

Yirgou, D., and Bradbury, J. F. 1968. Bacterial wilt of enset (*Ensete ventricosum*) incited by *Xanthomonas musacearum* sp. Phytopathology. 58:111–112.

Young, J. M., Bull, C. T., De Boer, S. H., Firrao, G., Saddler, G. E., Stead, D. E., et al. 2004. Names of Plant Pathogenic Bacteria, 1864 – 2004. Rev. Plant Pathol. 75:721–763.

Young, J. M., Dye, D. W., Bradbury, J. F., Panagopoulos, C. G., and Robbs, C. F. 1978. A proposed nomenclature and classification for plant pathogenic bacteria. New Zeal. J. Agric. Res. 21:153–177.

