## Supplementary Figure S1. Pathogenicity on banana for "Transfer of Xanthomonas campestris pv. arecae, and Xanthomonas campestris pv. musacearum to Xanthomonas vasicola (Vauterin) as Xanthomonas vasicola pv. arecae comb. nov., and Xanthomonas vasicola pv. musacearum comb. nov. and description of Xanthomonas vasicola pv. vasculorum pv. nov"

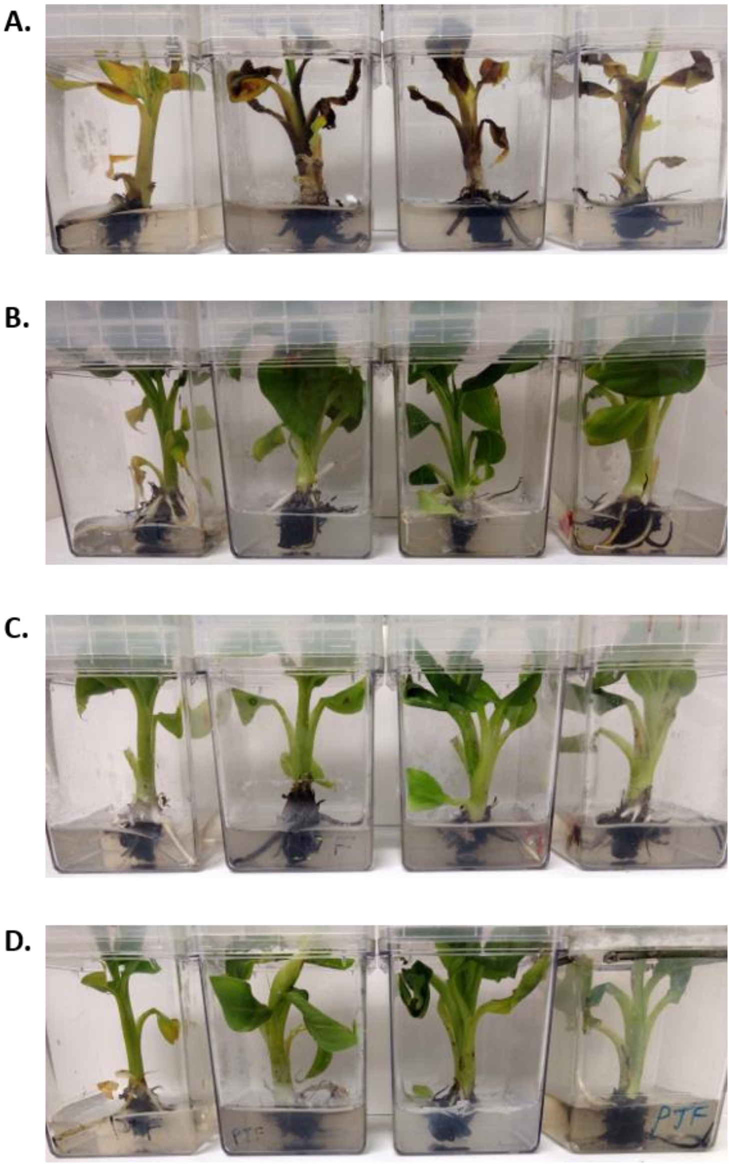


**Supplementary Figure S1. Pathogenicity tests of *Xanthomonas vasicola* strains on *Musa acuminata* (AAA Group) 'Grand Nain'.** Grand Nain banana plants in tissue culture 20 days post syringe inoculation at OD 0.2 with **A.** Xvm NCPPB 4433, **B.** 10 mM $MgCl_2$ , **C.** Xvv SAM119, **D.** Xvv NCPPB 702
